# A Deep Quantitative Proteome Turnover Platform for Human iPSC-derived Neurons

**DOI:** 10.64898/2026.03.14.711828

**Authors:** Ashley M. Frankenfield, Jamison Shih, Tao Zhang, Jiawei Ni, Wan Nur Atiqah binti Mazli, Edwin Lo, Yansheng Liu, Jiou Wang, Ling Hao

## Abstract

Quantitative evaluation of protein turnover in human neurons is crucial for understanding neuron homeostasis and guiding drug development for neurological diseases. However, measuring protein turnover in postmitotic neurons remains challenging due to the high dynamic range of protein half-lives and limited proteome coverage in SILAC (Stable Isotope Labeling by Amino acids in Cell culture) experiments. Despite broad applications of dynamic SILAC proteomics to measure protein turnover in rodent tissues and primary neurons, few studies have measured protein half-lives in human neurons with limited proteome coverage. Here, we established a comprehensive platform to quantify protein half-lives in human induced pluripotent stem cell (iPSC)-derived neurons. By integrating optimized dynamic SILAC labeling in human neuron cultures, extensive peptide fractionation, optimized data-dependent and data-independent LC-MS/MS acquisition methods, and a streamlined computational pipeline, we achieved deep and accurate measurement of 10,792 protein half-lives from 162,854 unique peptides. We then compared the protein turnover and abundances in iPSC-derived glutamatergic cortical neurons and spinal motor neurons, revealing globally conserved proteome dynamics alongside subtype-specific differences consistent with specialized neuronal functions. To enable broad community access, we created NeuronProfile (www.neuronprofile.com), an interactive web platform for exploring protein turnover, abundance, and subcellular location in human neurons. Together, this work provides a comprehensive analytical platform to assess human neuronal proteostasis and a foundational resource for neurological disease research and therapeutic development.

## Introduction

Neurons are highly specialized, long-lived cells that transmit and receive electrochemical signals throughout the nervous system^1^. As postmitotic cells that no longer divide, neurons are particularly susceptible to proteotoxic burden and metabolic stress. Impairments in protein degradation pathways and the accumulation of misfolded and aggregated proteins are strongly linked to the pathogenesis of numerous brain diseases^2,3^. Therefore, defining protein turnover in neurons is essential for understanding proteostasis in the central nervous system (CNS) and the mechanisms underlying neurological disorders. The development of CNS therapeutics remains exceptionally challenging. Knowledge of protein turnover is fundamental to rational drug development, as protein turnover rates directly influence target engagement duration, dosing frequency, and the design of targeted protein degradation strategies^4^. Protein half-lives also shape pharmacodynamic responses and biomarker changes during treatment^5^. Thus, a quantitative understanding of neuronal protein turnover is critical not only for elucidating fundamental neuronal homeostasis but also for guiding therapeutic development for neurological disorders.

Protein turnover reflects the dynamic balance between protein synthesis and degradation, which together determine the lifetime of proteins within a cell^6^. Protein turnover rates and half-lives can be measured by dSILAC (Dynamic Stable Isotope Labeling by Amino acids in Cell culture) proteomics, a quantitative approach that has been broadly applied across diverse cellular and animal systems^7–11^. By switching the nutrient source to the one containing stable heavy isotope-labeled amino acids, newly synthesized proteins incorporate heavy amino acids, while the abundances of light-labeled proteins decline due to degradation. Quantifying the heavy-to-light protein/peptide ratios at various time points allows the construction of protein/peptide degradation and synthesis curves to determine protein half-lives. However, substantial technical challenges remain for quantifying protein turnover. These include the wide dynamic range of protein half-lives, especially in postmitotic cells; missing values across isotope channels and time points; complex data modeling procedures; inconsistencies between peptide- and protein-level measurements; and limited proteome coverage due to increased MS spectral complexity in SILAC data.

Neuronal protein turnover has largely been characterized in primary rodent neurons and mouse tissues^9–16^. These studies have advanced our understanding of proteostasis dynamics, but species differences may limit the translation of these findings to human neurons. Recent advances in human induced pluripotent stem cells (iPSCs) technology have enabled the generation of functional human neurons *in vitro*^17^. Human somatic cells, such as dermal fibroblasts, can be reprogrammed into iPSCs and subsequently differentiated into neurons using defined small-molecule cocktail treatments or overexpression of lineage-specific transcription factors^18–22^. In our previous study, we measured protein half-lives in human iPSC-derived neurons for the first time and demonstrated that progranulin deficiency led to substantial alteration in protein turnover and impaired lysosomal dynamics^23^. However, proteome coverage was limited to fewer than 4,000 protein half-lives. Cavarischia-Rega *et al*. reported approximately 4300 protein half-lives in iPSC-derived dopaminergic neurons and further examined local protein synthesis and trafficking between axons and the soma^24^. Many drug targets and signaling regulators reside in the low-abundant range of the neuronal proteome, making them particularly difficult to detect and quantify with limited proteome coverage. Moreover, comparing protein turnover and expression across neuronal subtypes can illuminate how proteome dynamics support distinct functional programs, such as synaptic transmission, axonal maintenance, and neurotransmitter-specific signaling.

In this study, we established a comprehensive platform to measure global protein half-lives in human iPSC-derived neurons, including dSILAC labeling of iPSC-derived neurons, proteomics sample preparation with extensive offline LC fractionation, data-dependent and - independent acquisition (DDA & DIA) LC-MS/MS analysis, and a streamlined data analysis pipeline. Enabled by this new workflow, we measured the half-lives of 10,792 proteins and 162,854 unique peptides in human neurons. Comparing the protein half-life and abundances in glutamatergic cortical neurons and spinal motor neurons revealed a largely conserved global proteome landscape, alongside subtype-specific protein differences associated with specialized cortical and motor neuron functions. To enable broad access to our neuronal proteome resource, we created NeuronProfile (www.neuronprofile.com), an interactive web platform for visualizing and interrogating protein half-lives, abundances, and subcellular locations in human neurons.

## Experimental Section

### Human iPSC-derived Neuron Culture

Wild-type human iPSCs were differentiated into two types of neurons (cortical and motor neurons) using well-established protocols, as described previously^22,25^. Detailed steps for iPSC-neuron culture and differentiation are provided in **Supplemental Methods**. Glutamatergic cortical neurons and spinal motor neurons were matured in independent cultures and prepared for dynamic SILAC labeling by washing the cultures with warm phosphate-buffered saline (PBS) twice. Cultures were then switched to SILAC neuron medium (DMEM:F12 for SILAC medium, N2 Supplement, B-27 Supplement, NEAA, GlutaMAX, BDNF, GDNF, NT-3, laminin, doxycycline, and ^13^C_6_^15^N_2_ Lysine and/or ^13^C_6_ ^15^N_4_ Arginine). Neurons were harvested at 1, 2, 4, and 6 days (accurate to within 10 min) for multiple time point curve fitting. Neurons were gently washed with PBS twice and lysed in ice-cold lysis buffer (8M Urea, 150 mM sodium chloride, 50 mM Tris-HCl buffer).

### Dynamic SILAC Sample Preparation

Neuron protein sample preparation was conducted as described previously with minor adjustments^23,26^. Briefly, protein lysates were sonicated for 15 min in ice-cold water and clarified by centrifugation. Protein concentrations were determined using a DCA colorimetric assay (Bio-Rad). Protein lysates were reduced by 5 mM of Tris(2-carboxyethyl)phosphine (TCEP) and alkylated by 15 mM of iodoacetamide (IAA). Excessive IAA was quenched by dithiothreitol (DTT). Samples were diluted with 50 mM Tris-HCl buffer to get urea concentration below 1 M. Dynamic SILAC samples with Lys8 and Arg10 labels (Cambridge Isotope) were digested with Trypsin/Lys-C (Promega) at a 1:30 (enzyme: protein) ratio for 18 h at 37 °C. Samples with only Lys8 label were digested with Lys-C (Promega) for 18 h at 37 °C. Digestion was quenched by adding 10% trifluoroacetic acid (TFA) until the pH was lower than 2. Peptides were desalted using a Waters Oasis HLB plate and dried under a SpeedVac. All samples were stored at −30 °C until LC-MS analysis.

### Peptide Fractionation

Peptide samples were fractionated offline by either using a 96-well Waters HLB desalting plate or high pH reverse phase liquid chromatography. For HLB fractionation, the HLB plate was wet with HPLC-grade methanol and then equilibrated with water containing 1% TFA. Samples were loaded onto the plate and washed with 1% TFA for 2 times. Peptides were eluted in 4 subsequent fractions with 15%, 35%, 50%, and 99% MeOH with 1% TFA. Fractionated samples were dried down in a SpeedVac and stored at −30 °C. For offline LC fractionation, peptide samples were separated on a Waters XBridge BEH C18 column (3.5µm, 130 Å, 4.6 mm × 250 mm) connected to a Vanquish Flex UHPLC. Buffer A was 10 mM ammonium formate in water. Buffer B was 10 mM ammonium formate with 90% acetonitrile (ACN). Both buffers were adjusted to pH 10 using ammonium hydroxide. Column temperature was 50 °C and flow rate was 0.5 ml/min. LC gradient started from 5% B for 5 min, and then 5% B to 10% B in 2 min, 60% B in 35 min, 90% B for 10 min and equilibration at 5% B for 10 min. Fractions were collected every 60 s between 12 to 60 min using a Thermo Scientific Fraction Collector. All fractions were dried down and reconstituted in 0.1% formic acid (FA) in water before recombining the fractions based on LC-UV chromatogram.

### LC-MS/MS Analysis

All peptide samples were dried down, reconstituted in 0.1% formic acid (FA) in water, and clarified by centrifugation. LC-MS/MS analyses were conducted using a Dionex Ultimate 3000 RSLC system coupled with a Thermo Fisher Q-Exactive HF-X MS. LC buffer A was 0.1% FA in water, and buffer B was 0.1% FA in ACN. Samples were separated on an Easy-spray PepMAP RSLC C18 columns (2 µm, 100 Å, 75 µm) maintained at 55 °C. Single-shot and HLB fractionated samples were separated using a 3.5 h LC gradient on 75 cm and 50 cm columns, respectively. Offline LC fractionated samples were separated using a 100 min LC gradient on a 25 cm column. For data dependent acquisition (DDA), a precursor scan was taken from *m/z* 400 to 1000 with a resolving power of 120 K (at *m/z* 200 full width at half maximum (FWHM)). The automatic gain control (AGC) target was 1 X 10^6^ with a maximum injection time (MaxIT) of 60 ms. Precursors were isolated using a *m/z* 1.4 window using a Top 40 DDA method. A normalized collision energy (NCE) of 30% was used for fragmentation. MS/MS scans were acquired with a resolving power of 7.5 K, AGC of 2 X 10^5^, and MaxIT of 40 ms. For data independent acquisition (DIA), a survey scan was performed from *m/z* 400 to 1000 with a resolving power of 60K at *m/z* 200 FWHM, AGC of 1 X 10^6^, and MaxIT of 60 ms. Precursors were fragmented using an NCE of 30%. Fragments were detected using an *m/z* 8.0 staggered (overlapped) isolation pattern with 75 sequential DIA windows with a resolving power of 15K at 200 *m/z* FWHM, MaxIT of 40 ms, and NCE of 30%.

### Proteomics Data Analysis

DDA and DIA dynamic SILAC data were analyzed using MaxQuant (1.6.14)^27^ and Spectronaut (17.4) software, respectively. Swiss-Prot *Homo sapiens* database (reviewed) and our custom neuron-specific contaminant FASTA library (https://github.com/HaoGroup-ProtContLib) were used for protein/peptide identifications with a false discovery rate cutoff of 1%^28^. Trypsin/P or LysC enzyme was selected with a maximum of two missed cleavages. A fixed modification of cysteine carbamidomethylation and variable modifications of methionine oxidation and protein N-terminus acetylation were included. Precursor and fragment tolerance were set to 20 ppm. For MaxQuant, a standard multiplicity with two labels was used for data analysis. For Spectronaut, a labeled workflow was used with disabled precursor imputation and enabled in-silico generation of missing channels. Lys8 or Lys8/Arg10 was selected as the heavy channel. For DIA, b-ions were excluded during quantification to avoid ratio distortion. Peptide-level export files were used for downstream analysis using Python.

To calculate protein half-lives, reverse (“Rev_”) and contaminant proteins (“Cont_”) were first removed from all export files. To account for the QE-HFX orbitrap’s instruments limit of detection, intensities below 1000 and outlier heavy-to-light ratios (0.02 < Ratio < 100) were removed^29^. For fractionated samples, heavy-to-light ratios were averaged across all fractions. Relative isotope abundance (RIA) was calculated using the equation 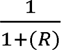, where R is the heavy-to-light ratio. Peptide RIAs from multiple time points were fitted to an exponential decay model with first-order kinetics: *RIA = e^-klOSSt^*, where t is the time point after medium switch, and k_loss_ is the rate of peptide decay. For postmitotic neurons, peptide half-life was then calculated as: *t_1/2_* = In(2)/*K_loss_*. Half-lives quantified with fewer than three time points or with an R^2^ < 0.8 were removed from the dataset. Peptide half-lives can also be estimated using a single time point equation assuming a steady state: 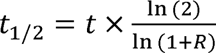. Each protein half-life was calculated as the harmonic mean of all unique peptides belonging to the protein.

### Bioinformatics Analyses

Statistical analyses were performed in R. Spearman’s correlation was calculated using the R function “stat_cor()”, and colored scatterplots were visualized using “ggplot2”. Subcellular locations were assigned to protein groups using Gene Ontology (GO) enrichment of cellular compartments from UniProt. Volcano plots were generated using VolcaNoseR^30^ and statistical significance was determined using a two-tailed Student’s *t-*test. Human Protein Atlas website was used to group proteins into classes, enzyme types, and ion channel types^31^. Cryo-electron microscopy structures publicly available at RCSB Protein Data Bank (PDB) were used to visualize protein complex structures^32^. Additionally, heatmaps were generated using the “pheatmap” package in R with Euclidean clustering^33^.

### Fluorescence Imaging

Neurons were cultured on PLO-coated 96-well plates for fluorescence imaging. Cortical neurons with stably integrated green fluorescence protein in the cytoplasm were imaged under a CELENA® S Digital inverted fluorescence microscope. Motor neurons were stained with 3 μM Calcein-AM (Invitrogen) and imaged under a Nikon Eclipse TS100 fluorescence microscope. Fluorescence images were processed using the Image-J software^34^.

## Results and Discussion

### Establishing a Comprehensive Workflow to Measure Protein Half-lives in Human iPSC-derived Neurons

Measuring protein turnover in postmitotic neurons is challenging due to the high dynamic range of protein half-lives (from minutes to months) and limited proteome coverage in SILAC experiments. To address these challenges, we established a comprehensive workflow and systematically optimized the experimental and data analysis parameters at each step. Human iPSC-derived neurons were cultured in normal light amino acid-containing medium until maturation and then switched to heavy amino acid-containing medium. Cultures were harvested at multiple timepoints (1, 2, 4, 6 days), followed by bottom-up proteomics (**Figure 1A**). For accurate and confident measurement of protein half-lives, we designed a data analysis pipeline including a series of filtering criteria, data modeling, and protein half-life calculations, described in detail in the **Experimental Section** and illustrated in **Figure 1B**. To remove contaminant proteins, we created a neuron culture-specific contaminant protein library based on the neuron medium composition and common contaminant proteins (**Supplementary Table S1**, available to download at https://github.com/HaoGroup-ProtContLib). An optimal set of data analysis parameters was selected for DDA and DIA data based on our recent study that benchmarked various SILAC proteomics software platforms^35^. Different peptide fractionation methods were evaluated to improve peptide separation and reduce LC-MS spectral complexity caused by the presence of both heavy and light peaks. Offline-LC fractionation provided significantly more protein/peptide IDs and more unique peptides per protein compared to the HLB solid phase extraction (SPE) fractionation and the single-shot no fractionation method (**Figure 1C**, **1D**, **1E**). A comparison of DDA and DIA LC-MS/MS methods revealed that the DIA approach quantified significantly more proteins and peptides compared to DDA (**Figure 1F**). DIA data presented fewer missing values for both heavy/light channels and replicates, as well as better reproducibility among technical replicates compared to DDA (**Figure 1G, Supplementary Figure S1A**). Peptide H/L ratios were consistent between DIA and DDA methods, indicating reliable quantification in both methods (**Figure 1H**). The single time point method, especially at 4 days, provided protein half-life measurements that were consistent with those obtained from the multiple time point curve-fitting approach (**Supplementary Figure S1B**). In summary, we presented a comprehensive and systematically optimized workflow, including dSILAC labeling of iPSC-derived neurons, proteomics sample preparation with extensive offline LC fractionation, LC-MS/MS analysis, and a streamlined data analysis pipeline, enabling us to measure global protein turnover in human neurons with state-of-the-art analytical depth (10,792 proteins and 162,854 unique peptides) and accuracy.

**Figure 1:**
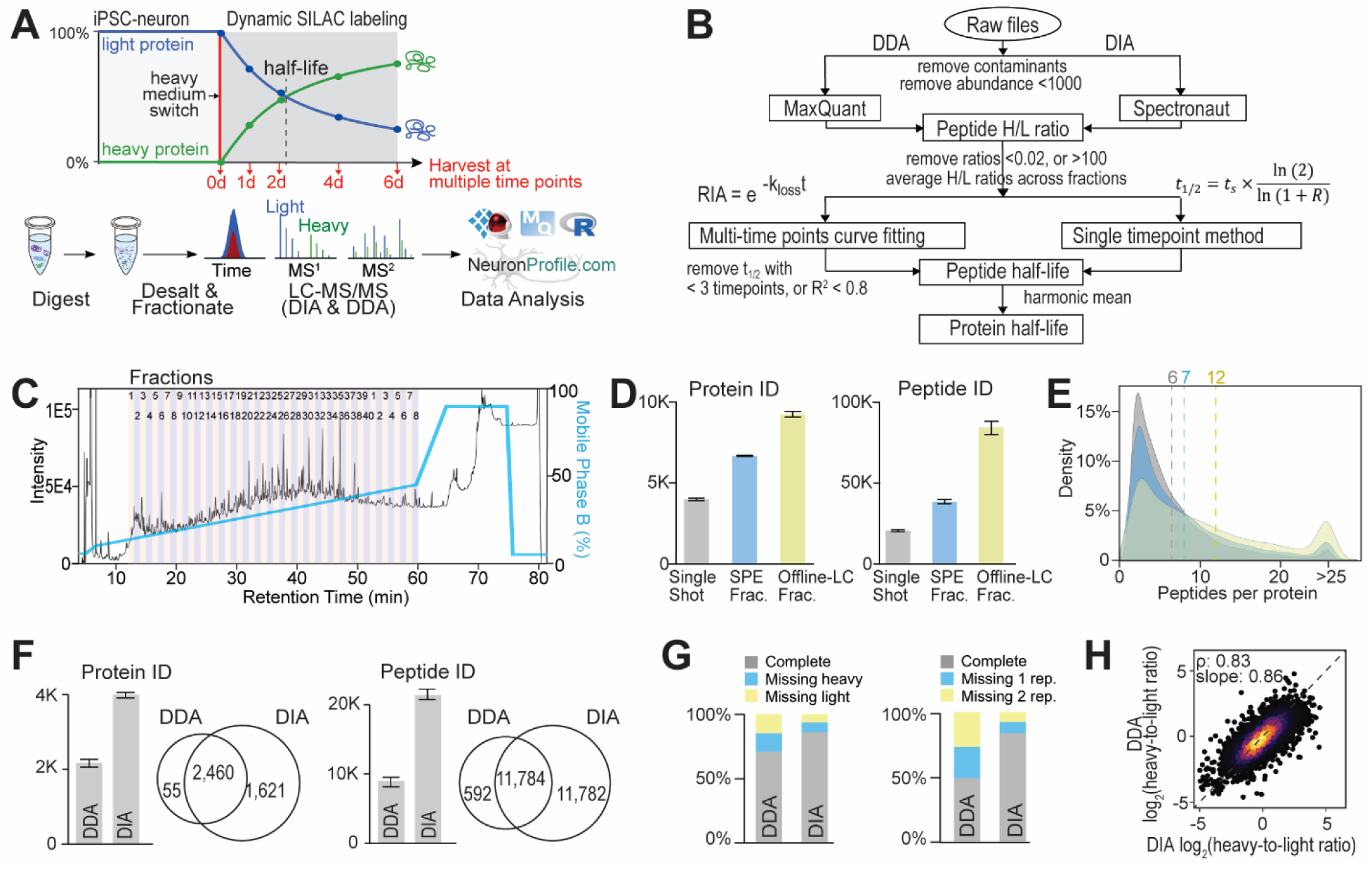
Establishing a comprehensive workflow for the deep and accurate measurement of protein half-lives in human iPSC-derived neurons. (A) Schematic overview to measure protein half-lives in human iPSC-derived neurons. (B) Data analysis flowchart for dynamic SILAC proteomics. (C) LC-UV/vis chromatogram of extensive offline-LC fractionation for peptide samples. (D) Comparison of the number of proteins and peptides quantified with single-shot, solid phase extraction fractionation (SPE Frac.) and offline-LC fractionation. (E) Distribution of the number of quantified unique peptides per protein from three methods. Dashed lines denote the average numbers. (F) Comparison of proteins and peptides quantified in DDA vs. DIA analysis of a 4-day single time point dSILAC neuron sample using the single-shot method. (G) Comparison of missing value percentages in heavy/light channel (left) and replicates (right) in DDA vs. DIA data. (H) Scatterplot correlation of heavy-to-light (H/L) ratios determined by DDA and DIA methods. ρ indicates Spearman’s correlation.

### Global Protein Turnover is Generally Conserved in Human iPSC-derived Motor and Cortical Neurons

To evaluate protein turnover in different neuron subtypes, we conducted dSILAC proteomics experiments in both human iPSC-derived glutamatergic cortical neurons and spinal motor neurons (**Figure 2A**). Glial cell marker GFAP was not identified in all samples, indicating the high purity of iPSC-derived neurons without glial cells. Despite differences in neuronal subtype and differentiation process, we found that global neuron protein turnover was generally conserved with similar half-life distributions and good correlation between cortical neurons (median of 4.0 d) and motor neurons (median of 4.2 d) (**Figure 2B, 2C**). Grouping proteins based on protein class and enzyme type showed similar overall protein half-lives in neuron subtypes (**Supplementary Figure S2**). Evaluating different half-life segments from both cortical and motor neuron proteins showed specifically enriched subcellular organelles (**Figure 2D**). E3 ubiquitin ligases (*e.g.* PJA1, PJA2, AMFR, RLIM, ZNRF1) and microtubule regulators (*e.g.* stathmins, kinesins) were among the fastest turnover proteins (half-life <1 day) in both cortical and motor neurons. Synaptic proteins (*e.g.* syntaxins, cadherins), Golgi membrane proteins, and SNARE complexes also showed relatively fast turnover rates. Mitochondrial inner membrane complexes and mitochondrial ribosome proteins had relatively slower turnover rates, consistent with previously reported studies on long-lived mitochondrial proteins^36^. Nucleosome (histones) and chromatin (HMGAs) proteins were among the slowest turnover proteins (half-life >10 days), consistent with other protein turnover studies^12^.

**Figure 2:**
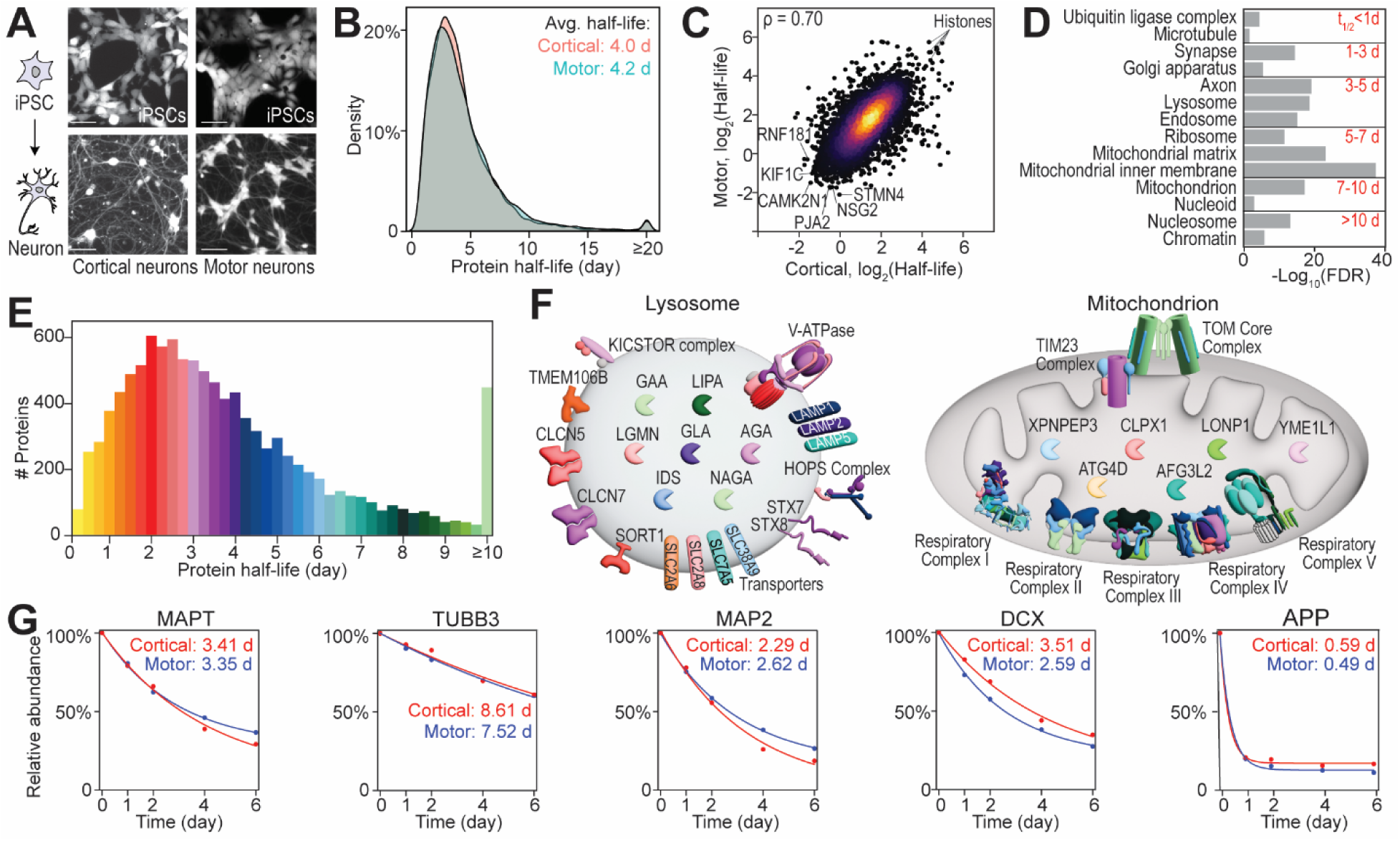
Global protein turnover in human iPSC-derived cortical and motor neurons. (A) Fluorescence microscopy imaging showing iPSCs and iPSC-derived cortical and motor neuron cultures. Scale bar is 50 µm. (B) Protein half-life distribution in cortical and motor neurons. (C) Scatterplot correlation of protein half-lives calculated from cortical and motor neurons. ρ indicates Spearman’s correlation. (D) GO-enrichment analysis showing significantly enriched subcellular locations of proteins with different half-life segments. (E) Average neuronal protein half-life distribution with color codes. (F) Example schematics of the lysosome and mitochondrion showing key proteins and protein complexes with color-coded half-lives. (G) Example protein degradation curves of key neuron markers.

Interestingly, although protein half-life distribution showed an organelle-dependent trend, proteins belonging to the same organelle can have a wide range of half-lives. Example lysosomal and mitochondrial proteins (color-coded half-lives based on **Figure 2E**) are shown in **Figure 2F**. Proteins belonging to the same complex tend to have similar half-lives^15^. For example, subunits from the v-ATPase complex (responsible for lysosome acidification^37^) had tightly clustered half-lives between 2 to 3.5 days, and proteins from the mitochondrial OXPHOS complexes (responsible for ATP production^38^) had longer half-lives over 6 days. We further examined the molecular protein functions with different ranges of half-lives as shown in **Supplementary Figure S3**. Short-lived proteins were involved in transcription regulation and ubiquitination processes. Long-lived proteins are involved in DNA binding and nucleosome binding. Key neuronal markers were highly expressed with very similar half-lives in cortical and motor neurons, such as tau (MAPT), TUJ1 (TUBB3), MAP2, DCX, APP, NeuN (RBFOX3), and NCAM1, indicating the maturation of both neuron subtypes and consistent core neuronal functions (**Figure 2G**).

### Human iPSC-derived Motor and Cortical Neurons Exhibit Differential Protein Turnover and Expression Related to Neuron Subtype Functions

With an overall comparable global protein turnover between neuron subtypes, we further investigated if any proteins exhibit significantly different half-lives due to subtype-specific neuronal functions. We first ranked the protein half-life ratios from motor/cortical neurons as shown in **Figure 3A**. Proteins related to axonal fasciculation, neuron recognition, and synapse organization showed shorter half-lives in motor neurons (**Figure 3B**). Faster turnover of axonal proteins may help spinal motor neurons develop long axons and bundle together to travel long distances to connect with target muscles^39^. On the other hand, proteins involved in glycosylation and metabolism showed faster turnover in cortical neurons, which are excitatory interneurons that secrete glutamate and consume more than 70% of the total neuronal metabolism^40^. Proteins from different ion channels showed differences between cortical and motor neurons (**Figure 3C**). Voltage-gated sodium channel proteins (*e.g.* SCN9A, SCN2A, SCN3A, SCN3B), which mediate action potential initiation and propagation, were more stable in cortical neurons, as exemplified in **Figure 3D** for SCN9A^41^. Sodium-dependent phosphate transporters SLC20A1 and SLC20A2, which have been linked to neuronal plasticity and cognition, were more stable in cortical neurons^42^. Potassium voltage-gated channel proteins (KCNA6, KCND1, KCNH2, KCNC1, KCND2, KCND3, KCNB2), which have been linked to motor neuron diseases, were more stable in motor neurons^43^.

**Figure 3:**
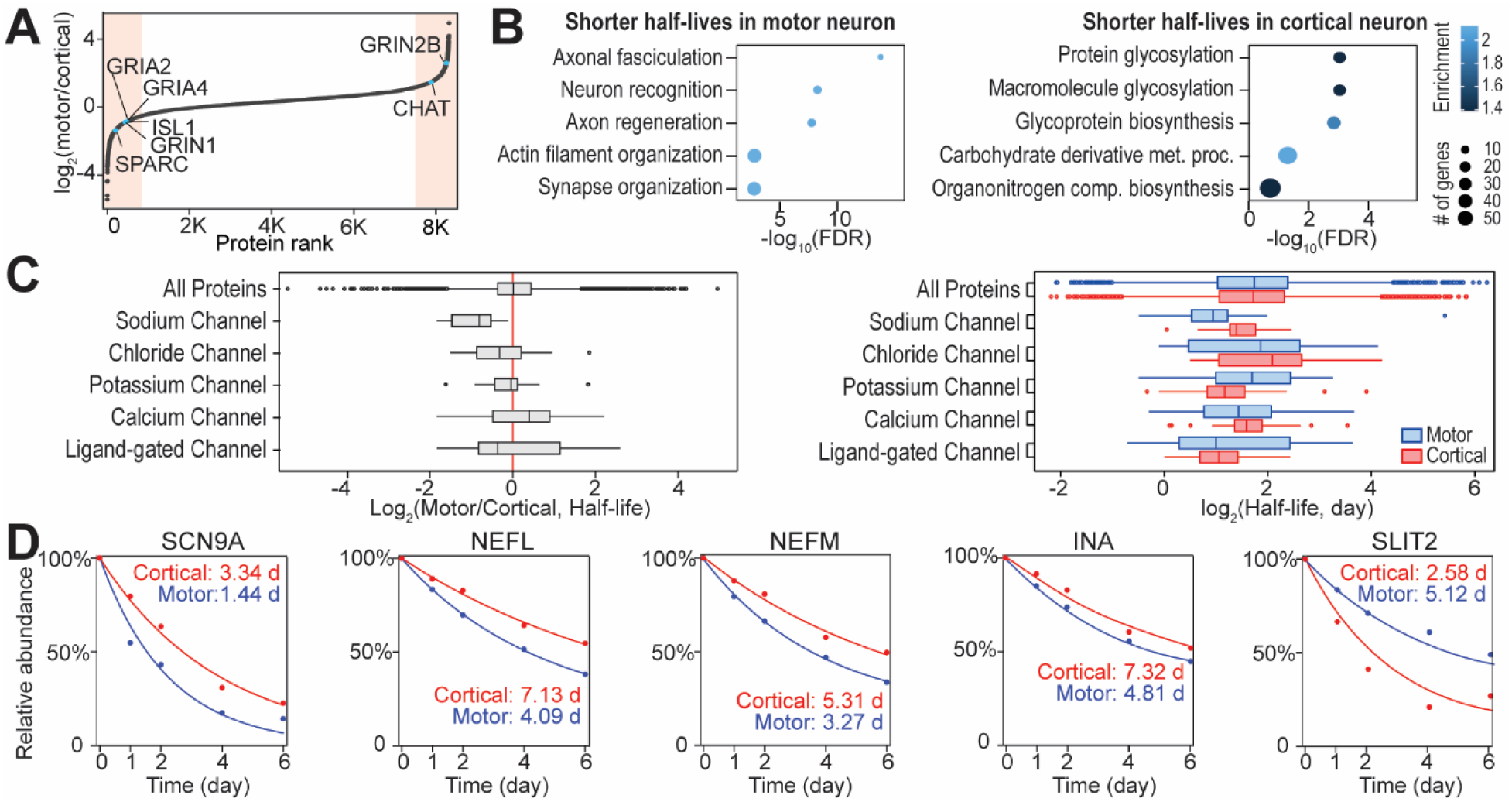
Comparing protein half-lives in human iPSC-derived cortical and motor neurons. (A) Ranking scatter plot showing the ratios of protein half-lives measured in motor and cortical neurons. (B) GO enrichment analysis showing biological processes enriched from the top 5% of proteins with significantly different half-lives in motor and cortical neurons. (C) Box plots of half-life ratios (left) and distributions (right) from proteins belonging to different ion channels. (D) Example protein degradation curves with different half-lives in cortical and motor neurons.

To further characterize proteomic differences between the neuronal subtypes, we compared protein abundances in motor vs. cortical neurons by label-free DIA proteomics and assessed the relationship between protein expression and turnover. Overall protein abundances were highly correlated between neuronal subtypes (**Figure 4A**). However, many proteins showed differential expressions between motor and cortical neurons (**Figure 4B**). Cortical glutamatergic neuron markers PRSS12 and NEUROG2 were exclusively detected in cortical neurons with a half-life of 4.2 and 0.4 days, respectively. NEUROG2 transcription factor is overexpressed to drive cortical neuron differentiation in the i^3^Neuron^21^ cell line and was only detected in cortical neurons. Group 3 metabotropic glutamate receptors (GRM4, GRM7, GRM8; half-life ∼2.7 days) and neuron-specific glutamate transporters (SLC17A6, SLC17A7) were only detected in cortical glutamatergic neurons. On the other hand, HOX transcription factors (HoxC8, HoxC6, HoxA7, etc.), responsible for the terminal differentiation of cholinergic and motor neurons, were only detected in motor neurons^44^. GO-enrichment analysis showed that proteins related to metabolic processes were more abundant in motor neurons, while proteins related to secretion and synaptic vesicles were more abundant in cortical neurons **(Figure 4B**, **4C)**. Extracellular chemorepellents and their receptors (SLIT1, SLIT2, UNC5C) responsible for blocking the formation of axons in incorrect positions, were more abundant in motor neurons with long axons^45^. Furthermore, proteins involved in glutamate biosynthesis (ASS1, GLS), neurotransmitter secretion (SCG2, SCG3, PCLO), and cell-cell adhesion (ADGRL3, ADGRL2) were significantly more abundant in the cortical neurons^46,47^.

**Figure 4:**
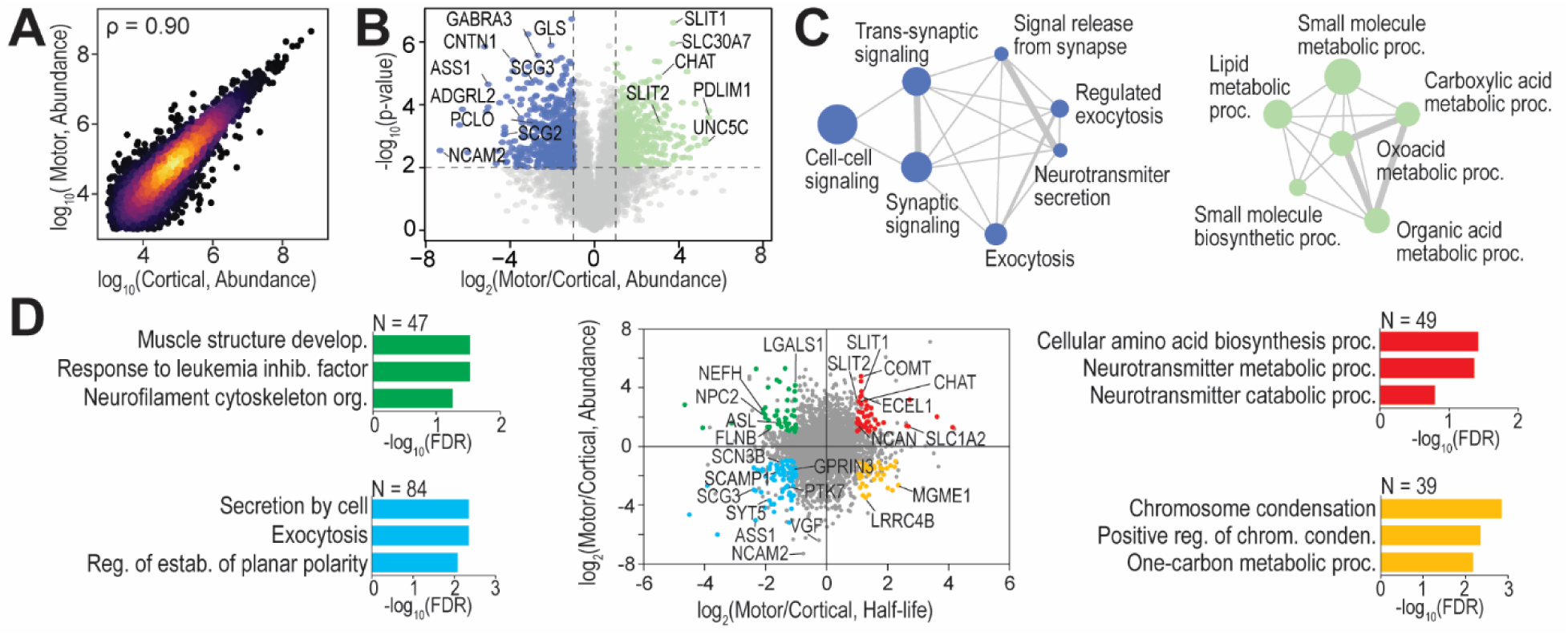
Comparing protein abundances and the relationship with half-lives in human iPSC-derived cortical and motor neurons. (A) Scatterplot correlation of protein abundances measured from cortical and motor neurons. ρ indicates Spearman’s correlation. (B) Volcano plot of protein abundance comparison. Dashed lines denote p-value of 0.01 and fold change of 2. (C) GO enrichment analysis showing biological processes enriched from proteins with significantly higher abundances in cortical neurons (left) and motor neurons (right). (D) Scatter plot showing the relationship between protein half-lives and protein abundance comparisons of motor vs. cortical neurons. GO-enrichment analysis showing biological processes enriched from different segments of the figure with significant differences.

The relationship between protein half-life and expression is largely unknown, especially in neurons. To further understand their relationship, we plotted the motor/cortical protein ratios of abundance and half-life and highlighted significantly changed proteins (p-value <0.05, fold change >2) and their enriched GO-terms in **Figure 4D**. Proteins involved in neurofilament and muscle structure development presented faster turnover and higher abundance in motor neurons (upper left). Proteins related to secretion and exocytosis showed faster turnover and lower abundance in motor neurons (lower left). Proteins related to neurotransmitter and metabolic processes showed faster turnover and lower abundance in cortical neurons (upper right). Proteins related to chromosome condensation showed faster turnover and higher abundance in cortical neurons (lower right). Proteins specific to motor and cortical neuron functions showed interesting differences in protein turnover and expression. For example, the cholinergic neuron marker, choline acetyltransferase (CHAT), was 8-fold more abundant in cholinergic motor neurons (*p*<0.0001) and also more stable (t_1/2_=4.6 d) compared to glutamatergic cortical neurons (t_1/2_=1.7 d). The axon initiation segment marker ankyrinG (ANK3), which is critical for maintaining neuron polarity, was 1.9-fold more abundant and less stable in motor neurons^48^. Synaptotagmins (SYT1, SYT2, SYT5, SYT7), major calcium sensors for neurotransmitter release and exocytosis, were more abundant and more stable in cortical neurons^49^. Neurofilament proteins were generally more abundant and stable in cortical neurons. In contrast, the heavy chain NEFH was markedly enriched in motor neurons (fold change = 5.6, *p* = 0.0019) and is a key determinant of axonal diameter^50^. All neurofilament proteins—including NEFL, NEFM, NEFH, INA, and PRPH—exhibited faster turnover in motor neurons. These proteins provide structural support for axons and serve as biomarkers for motor neuron diseases such as Amyotrophic Lateral Sclerosis (ALS)^50^.

### Cross-System Comparison of Protein Half-lives in Neural Models

Various systems have been used to model the human brain and measure neuronal protein turnover, such as rodent brains, primary neuron culture from rodents, and human iPSC-derived neurons^9,10,12–14^. To understand how model choice affects protein half-life measurement, we compared protein half-lives measured in our human iPSC-derived neurons with previously published datasets from iPSC-derived dopaminergic neurons, mouse and rat primary neurons, and mouse brain cortex tissues. Our dataset quantified the largest number of protein half-lives, providing the half-lives of 3,896 unique proteins that were not measured in previous studies (**Figure 5A**). Protein half-life results were clustered based on model systems. Interestingly, primary mouse neurons were clustered together with iPSC-derived neurons, while primary rat neurons were more closely clustered with mouse tissues **(Figure 5A, 5B)**. We then further examined synaptic proteins from both presynaptic and postsynaptic compartments (**Figure 5C, Supplementary Figure S4**). Synaptic proteins showed similar turnover in neuron cultures but had longer half-lives in tissue samples, likely due to more advanced developmental stage of tissue samples *in vivo*^14,51^. For example, proteins involved with structural maintenance of the synapse (DLGAPs, DLGs, HOMERs) had significantly longer half-lives in tissues compared to cell culture models. Protein half-lives calculations and data analysis pipelines used in tissue analysis can also contribute to these differences^52^.

**Figure 5:**
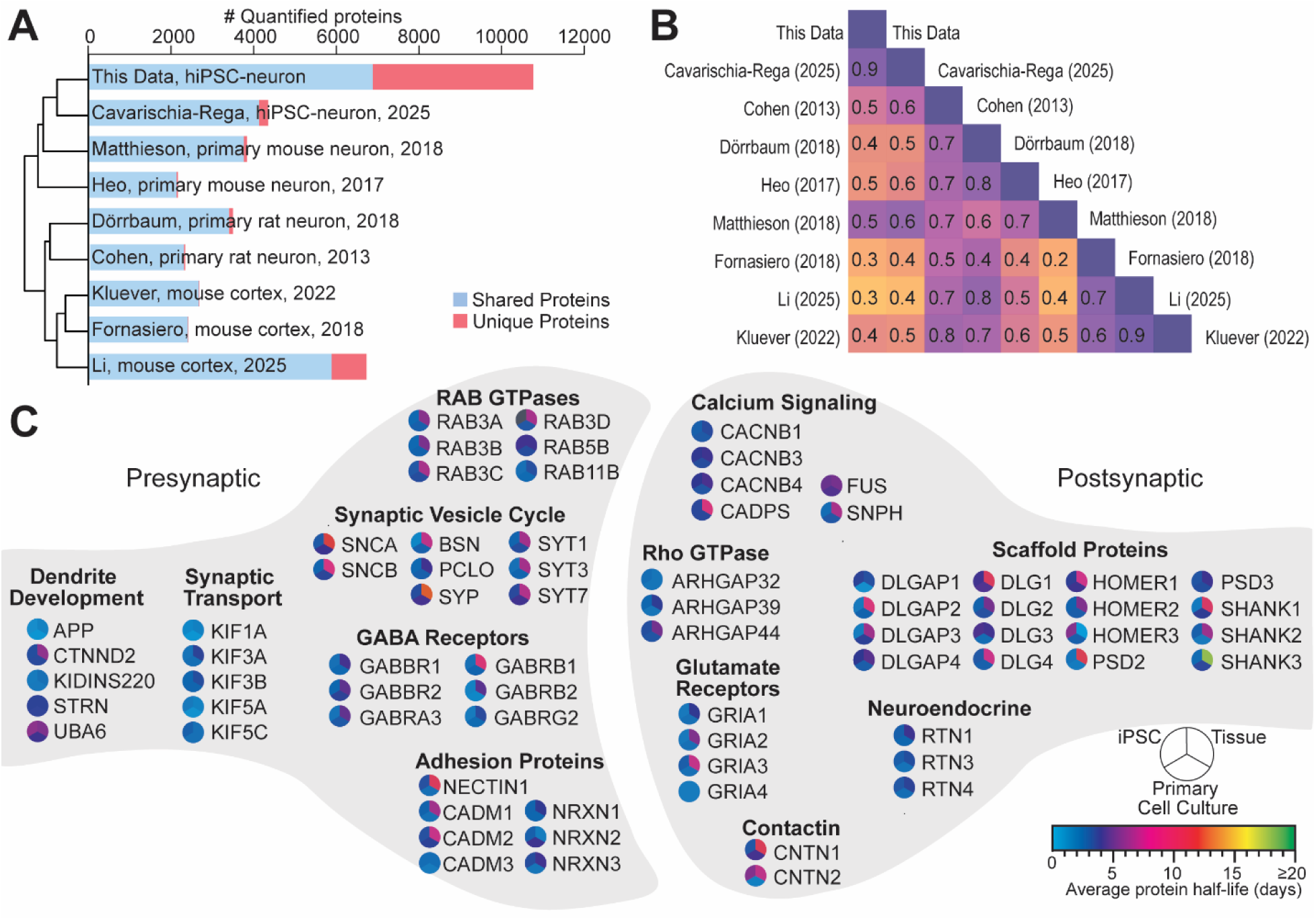
Evaluating protein turnover in iPSC-derived neuron culture, primary rodent neuron culture, and mouse brain tissues. (A) Bar chart comparing the number of protein half-lives measured in our human iPSC-neuron dataset and previously published datasets from rodent primary neuron and brain tissues. Hierarchical clustering using protein half-lives from different studies is shown on the left. (B) Correlation of protein half-lives from different datasets. Numbers indicate ρ from Spearman’s correlation. (C) Schematic of average synaptic protein half-lives measured in different sample types.

### NeuronProfile: An Interactive Resource for Protein Half-life and Abundance in Human iPSC-derived Neurons

Understanding neuronal protein stability is essential for predicting pharmacodynamic responses, interpreting biomarker dynamics, and anticipating protein accumulation in CNS drug development. To support the accessibility and utility of our comprehensive neuron protein datasets, we developed NeuronProfile (www.neuronprofile.com), an interactive website that enables users to conveniently visualize, query, and analyze protein half-lives, abundances, and subcellular localizations in human iPSC-derived neurons (**Figure 6A**). The platform features an interactive scatter plot of protein half-lives (**Figure 6B**) and displays degradation and synthesis curves at both the protein and peptide levels, as illustrated for alpha-synuclein (SNCA) in **Figure 6C**. In addition, we estimated protein copy numbers per cell using the “proteomic ruler” approach^53^. Users can systematically search and compare protein copy numbers and half-lives across motor and cortical neuron types, making our NeuronProfile website a valuable resource to support hypothesis generation, mechanistic studies, and drug development in CNS research.

**Figure 6:**
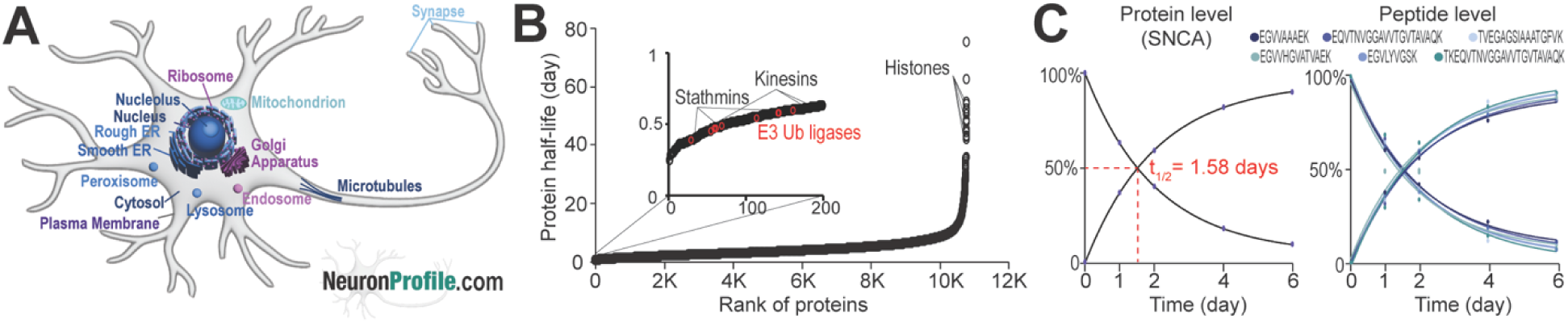
Establishing an interactive website (NeuronProfile.com) to visualize protein turnover and abundances with subcellular locations in human neurons. (A) Schematic of neurons showing color-coded average protein half-lives from different organelles. Protein half-life distribution with color codes is shown on the website and in Figure 3E. (B) Scatter plot distribution of 10,972 protein half-lives measured from human iPSC-derived neurons. (C) Example degradation and synthesis curves of alpha-synuclein (SNCA) protein and peptides.

### Conclusions

Measuring protein half-lives in neurons is fundamental to understanding CNS proteostasis and guiding rational development of therapeutics targeting neuronal proteins in brain diseases. In this study, we established a deep dynamic SILAC workflow in human iPSC-derived neurons, enabling the quantification of half-lives and abundance for over 10K neuronal proteins and 150K unique peptides. Comparative analyses of glutamatergic cortical neurons and spinal motor neurons revealed that global protein turnover is largely conserved between neuron subtypes, with a median half-life of approximately four days. Notably, proteins associated with subtype-specific neuronal functions exhibited distinct expression levels and turnover kinetics. To maximize the impact and accessibility of this resource, we created NeuronProfile (www.neuronprofile.com), an interactive platform for exploring protein turnover and abundance in human neurons. By sharing these deep neuronal proteome turnover and expression datasets with the scientific community, this work provides a valuable resource that enables cross-study comparisons, improves computational modeling of proteome dynamics, and supports the rational design of protein-targeted therapies for neurological diseases.

## Author Contributions

A.M.F. and L.H. designed the study. J.N. conducted iPSC-derived cortical neuron cell culture and fluorescence imaging. T.Z. conducted iPSC-derived motor neuron cell culture and imaging under the supervision of J.W. A.M.F, J.S., and W.M. conducted proteomic experiments, LC-MS analysis, and data analysis with suggestions from Y.L.. E.L developed the multiple-time point curve fitting Python code and source code for the NeuronProfile website. A.M.F wrote the first draft of the manuscript with revision from L.H. and Y.L. All authors have read and agreed to the final version of the manuscript.

## Data availability

All MS raw files have been deposited to ProteomeXchange via MassIVE with the project accession ID: PXD049255. The neuron-specific contaminant library is freely available at https://github.com/HaoGroup-ProtContLib. The proteomics results in this study can be visualized in an interactive format at https://neuronprofile.com.

## Acknowledgements

This study is supported by the NIH grants (R01NS121608 for L.H., R01NS074324 for J.W.). A.M.F acknowledges the Willard & Marilynn Sweetser Scholarship from the Metropolitan Chapter of Achievement Rewards for College Scientists (ARCS) Foundation. The authors would like to thank Tracey Lo for her feedback and assistance on the graphical design of NeuronProfile website.

## Supporting Information

- **Supplemental Methods**: Human iPSC-derived Neuron Culture.
- **Supplementary Figure S1:** Correlation analyses of DDA and DIA dynamic SILAC proteomics data from human iPSC-derived neurons.
- **Supplementary Figure S2:** Comparing protein half-lives from different protein classes in human motor and cortical neurons.
- **Supplementary Figure S3:** GO-enrichment analyses showing enriched molecular functions of proteins with different ranges of half-lives in human neurons.
- **Supplementary Figure S4:** Heatmap clustering of synaptic protein half-lives measured in our human neuron data and published datasets from rodent neurons and brain tissues.
- **Supplementary Table S1**: Neuron specific contaminant library.
- **Supplementary Table S2**: Protein turnover and abundance in human iPSC-derived cortical and motor neurons.

## References

(1) Levitan, I. B.; Kaczmarek, L. K. The Neuron; Oxford University Press, 2015. 10.1093/med/9780199773893.001.0001.

(2) Wilson, D. M.; Cookson, M. R.; Van Den Bosch, L.; Zetterberg, H.; Holtzman, D. M.; Dewachter, I. Hallmarks of Neurodegenerative Diseases. Cell 2023, 186 (4), 693–714. 10.1016/J.CELL.2022.12.032.

(3) Ross, C. A.; Poirier, M. A. Protein Aggregation and Neurodegenerative Disease. Nat. Med. 2004, 10 (S7), S10–S17. 10.1038/nm1066.

(4) Pangalos, M. N.; Schechter, L. E.; Hurko, O. Drug Development for CNS Disorders: Strategies for Balancing Risk and Reducing Attrition. Nat. Rev. Drug Discov. 2007, 6 (7), 521–532. 10.1038/nrd2094.

(5) Beller, N. C.; Wang, Y.; Hummon, A. B. Evaluating the Pharmacokinetics and Pharmacodynamics of Chemotherapeutics within a Spatial SILAC-Labeled Spheroid Model System. Anal. Chem. 2023, 95 (30), 11263–11272. 10.1021/acs.analchem.3c00905.

(6) Pratt, J. M.; Petty, J.; Riba-Garcia, I.; Robertson, D. H. L.; Gaskell, S. J.; Oliver, S. G.; Beynon, R. J. Dynamics of Protein Turnover, a Missing Dimension in Proteomics. Mol. Cell. Proteomics 2002, 1 (8), 579–591. 10.1074/mcp.M200046-MCP200.

(7) Doherty, M. K.; Hammond, D. E.; Clague, M. J.; Gaskell, S. J.; Beynon, R. J. Turnover of the Human Proteome: Determination of Protein Intracellular Stability by Dynamic SILAC. J. Proteome Res. 2009, 8 (1), 104–112. 10.1021/pr800641v.

(8) Cambridge, S. B.; Gnad, F.; Nguyen, C.; Bermejo, J. L.; Krüger, M.; Mann, M. Systems-Wide Proteomic Analysis in Mammalian Cells Reveals Conserved, Functional Protein Turnover. J. Proteome Res. 2011, 10 (12), 5275–5284. 10.1021/pr101183k.

(9) Mathieson, T.; Franken, H.; Kosinski, J.; Kurzawa, N.; Zinn, N.; Sweetman, G.; Poeckel, D.; Ratnu, V. S.; Schramm, M.; Becher, I.; Steidel, M.; Noh, K. M.; Bergamini, G.; Beck, M.; Bantscheff, M.; Savitski, M. M. Systematic Analysis of Protein Turnover in Primary Cells. Nat. Commun. 2018, 9 (1), 1–10. 10.1038/s41467-018-03106-1.

(10) Li, W.; Dasgupta, A.; Yang, K.; Wang, S.; Hemandhar-Kumar, N.; Yarbro, J. M.; Hu, Z.; Salovska, B.; Fornasiero, E. F.; Peng, J.; Liu, Y. An Extensive Atlas of Proteome and Phosphoproteome Turnover Across Mouse Tissues and Brain Regions. October 17, 2024. 10.1101/2024.10.15.618303.

(11) Yarbro, J. M.; Han, X.; Dasgupta, A.; Yang, K.; Liu, D.; Shrestha, H. K.; Zaman, M.; Wang, Z.; Yu, K.; Lee, D. G.; Vanderwall, D.; Niu, M.; Sun, H.; Xie, B.; Chen, P.-C.; Jiao, Y.; Zhang, X.; Wu, Z.; Chepyala, S. R.; Fu, Y.; Li, Y.; Yuan, Z.-F.; Wang, X.; Poudel, S.; Vagnerova, B.; He, Q.; Tang, A.; Ronaldson, P. T.; Chang, R.; Yu, G.; Liu, Y.; Peng, J. Human and Mouse Proteomics Reveals the Shared Pathways in Alzheimer’s Disease and Delayed Protein Turnover in the Amyloidome. Nat. Commun. 2025, 16 (1), 1533. 10.1038/s41467-025-56853-3.

(12) Savas, J. N.; Toyama, B. H.; Xu, T.; Yates, J. R.; Hetzer, M. W. Extremely Long-Lived Nuclear Pore Proteins in the Rat Brain. Science. American Association for the Advancement of Science February 24, 2012, p 942. 10.1126/science.1217421.

(13) Fornasiero, E. F.; Mandad, S.; Wildhagen, H.; Alevra, M.; Rammner, B.; Keihani, S.; Opazo, F.; Urban, I.; Ischebeck, T.; Sakib, M. S.; Fard, M. K.; Kirli, K.; Centeno, T. P.; Vidal, R. O.; Rahman, R. U.; Benito, E.; Fischer, A.; Dennerlein, S.; Rehling, P.; Feussner, I.; Bonn, S.; Simons, M.; Urlaub, H.; Rizzoli, S. O. Precisely Measured Protein Lifetimes in the Mouse Brain Reveal Differences across Tissues and Subcellular Fractions. Nat. Commun. 2018, 9 (1). 10.1038/s41467-018-06519-0.

(14) Dörrbaum, A. R.; Alvarez-Castelao, B.; Nassim-Assir, B.; Langer, J. D.; Schuman, E. M. Proteome Dynamics during Homeostatic Scaling in Cultured Neurons. Elife 2020, 9, 1–28. 10.7554/eLife.52939.

(15) Dörrbaum, A. R.; Kochen, L.; Langer, J. D.; Schuman, E. M. Local and Global Influences on Protein Turnover in Neurons and Glia. Elife 2018, 7, 1–24. 10.7554/eLife.34202.

(16) Basisty, N.; Meyer, J. G.; Schilling, B. Protein Turnover in Aging and Longevity. Proteomics 2018, 18 (5–6). 10.1002/pmic.201700108.

(17) Dolmetsch, R.; Geschwind, D. H. The Human Brain in a Dish: The Promise of IPSC-Derived Neurons. Cell 2011, 145 (6), 831–834. 10.1016/j.cell.2011.05.034.

(18) Karumbayaram, S.; Novitch, B. G.; Patterson, M.; Umbach, J. A.; Richter, L.; Lindgren, A.; Conway, A. E.; Clark, A. T.; Goldman, S. A.; Plath, K.; Wiedau-Pazos, M.; Kornblum, H. I.; Lowry, W. E. Directed Differentiation of Human-Induced Pluripotent Stem Cells Generates Active Motor Neurons. Stem Cells 2009, 27 (4), 806–811. 10.1002/stem.31.

(19) Lu, J.; Zhong, X.; Liu, H.; Hao, L.; Huang, C. T. L.; Sherafat, M. A.; Jones, J.; Ayala, M.; Li, L.; Zhang, S. C. Generation of Serotonin Neurons from Human Pluripotent Stem Cells. Nat. Biotechnol. 2016, 34 (1), 89–94. 10.1038/nbt.3435.

(20) Zhang, Y.; Pak, C. H.; Han, Y.; Ahlenius, H.; Zhang, Z.; Chanda, S.; Marro, S.; Patzke, C.; Acuna, C.; Covy, J.; Xu, W.; Yang, N.; Danko, T.; Chen, L.; Wernig, M.; Südhof, T. C. Rapid Single-Step Induction of Functional Neurons from Human Pluripotent Stem Cells. Neuron 2013, 78 (5), 785–798. 10.1016/j.neuron.2013.05.029.

(21) Wang, C.; Ward, M. E.; Chen, R.; Liu, K.; Tracy, T. E.; Chen, X.; Xie, M.; Sohn, P. D.; Ludwig, C.; Meyer-Franke, A.; Karch, C. M.; Ding, S.; Gan, L. Scalable Production of IPSC-Derived Human Neurons to Identify Tau-Lowering Compounds by High-Content Screening. Stem Cell Reports 2017, 9 (4), 1221–1233. 10.1016/j.stemcr.2017.08.019.

(22) Du, Z. W.; Chen, H.; Liu, H.; Lu, J.; Qian, K.; Huang, C. T. L.; Zhong, X.; Fan, F.; Zhang, S. C. Generation and Expansion of Highly Pure Motor Neuron Progenitors from Human Pluripotent Stem Cells. Nat. Commun. 2015, 6, 1–9. 10.1038/ncomms7626.

(23) Hasan, S.; Fernandopulle, M. S.; Humble, S. W.; Frankenfield, A. M.; Li, H.; Prestil, R.; Johnson, K. R.; Ryan, B. J.; Wade-Martins, R.; Ward, M. E.; Hao, L. Multi-Modal Proteomic Characterization of Lysosomal Function and Proteostasis in Progranulin-Deficient Neurons. Mol. Neurodegener. 2023, 18 (1), 87. 10.1186/s13024-023-00673-w.

(24) Cavarischia-Rega, C.; Sharma, K.; Fitzgerald, J. C.; Macek, B. Proteome Dynamics in IPSC-Derived Human Dopaminergic Neurons. Molecular and Cellular Proteomics 2024, 23 (10). 10.1016/j.mcpro.2024.100838.

(25) Fernandopulle, M. S.; Prestil, R.; Grunseich, C.; Wang, C.; Gan, L.; Ward, M. E. Transcription Factor–Mediated Differentiation of Human IPSCs into Neurons. Curr. Protoc. Cell Biol. 2018, 79 (1), 1–48. 10.1002/cpcb.51.

(26) Lee, G. Bin; Mazli, W. N. A. binti; Hao, L. Multiomics Evaluation of Human IPSCs and IPSC-Derived Neurons. J. Proteome Res. 2024, 23 (8), 3149–3160. 10.1021/acs.jproteome.3c00790.

(27) Cox, J.; Neuhauser, N.; Michalski, A.; Scheltema, R. A.; Olsen, J. V.; Mann, M. Andromeda: A Peptide Search Engine Integrated into the MaxQuant Environment. J. Proteome Res. 2011, 10 (4), 1794–1805. 10.1021/pr101065j.

(28) Frankenfield, A. M.; Ni, J.; Ahmed, M.; Hao, L. Protein Contaminants Matter: Building Universal Protein Contaminant Libraries for DDA and DIA Proteomics. J. Proteome Res. 2022, 21 (9), 2104–2113. 10.1021/acs.jproteome.2c00145.

(29) Frankenfield, A. M.; Yang, K. L.; Mazli, W. N. A. binti; Shih, J.; Yu, F.; Lo, E.; Nesvizhskii, A. I.; Hao, L. Benchmarking SILAC Proteomics Workflows and Data Analysis Platforms. Molecular & Cellular Proteomics 2025, 24 (6), 100980. 10.1016/j.mcpro.2025.100980.

(30) Goedhart, J.; Luijsterburg, M. S. VolcaNoseR Is a Web App for Creating, Exploring, Labeling and Sharing Volcano Plots. Sci. Rep. 2020, 10 (1). 10.1038/s41598-020-76603-3.

(31) Uhlén, M.; Fagerberg, L.; Hallström, B. M.; Lindskog, C.; Oksvold, P.; Mardinoglu, A.; Sivertsson, Å.; Kampf, C.; Sjöstedt, E.; Asplund, A.; Olsson, I.; Edlund, K.; Lundberg, E.; Navani, S.; Szigyarto, C. A.-K.; Odeberg, J.; Djureinovic, D.; Takanen, J. O.; Hober, S.; Alm, T.; Edqvist, P.-H.; Berling, H.; Tegel, H.; Mulder, J.; Rockberg, J.; Nilsson, P.; Schwenk, J. M.; Hamsten, M.; von Feilitzen, K.; Forsberg, M.; Persson, L.; Johansson, F.; Zwahlen, M.; von Heijne, G.; Nielsen, J.; Pontén, F. Tissue-Based Map of the Human Proteome. Science (1979). 2015, 347 (6220). 10.1126/science.1260419.

(32) Berman, H. M., Westbrook, J., Feng, Z., Gilliland, G., Bhat, T.N., Weissig, H., Shindyalov, I.N., Bourne, P. E. The Protein Data Bank. Nucleic Acids Res. 2000, 28 (1), 235–242. 10.1093/nar/28.1.235.

(33) McInnes, L.; Healy, J.; Saul, N.; Großberger, L. UMAP: Uniform Manifold Approximation and Projection. J. Open Source Softw. 2018, 3 (29), 861. 10.21105/joss.00861.

(34) Schneider, C. A.; Rasband, W. S.; Eliceiri, K. W. NIH Image to ImageJ: 25 Years of Image Analysis. Nat. Methods 2012, 9 (7), 671–675. 10.1038/nmeth.2089.

(35) Frankenfield, A. M.; Yang, K. L.; Mazli, W. N. A. binti; Shih, J.; Yu, F.; Lo, E.; Nesvizhskii, A. I.; Hao, L. Benchmarking SILAC Proteomics Workflows and Data Analysis Platforms. Molecular & Cellular Proteomics 2025, 24 (6), 100980. 10.1016/j.mcpro.2025.100980.

(36) Bomba-Warczak, E.; Edassery, S. L.; Hark, T. J.; Savas, J. N. Long-Lived Mitochondrial Cristae Proteins in Mouse Heart and Brain. Journal of Cell Biology 2021, 220 (9). 10.1083/jcb.202005193.

(37) Recchi, C.; Chavrier, P. V-ATPase: A Potential PH Sensor. Nat. Cell Biol. 2006, 8, 107–109. 10.1038/ncb0206-107.

(38) Senior, A. E. ATP Synthesis by Oxidative Phosphorylation. Physiol. Rev. 1988, 68 (1), 177–231. 10.1152/physrev.1988.68.1.177.

(39) Stifani, N. Motor Neurons and the Generation of Spinal Motor Neuron Diversity. Front. Cell. Neurosci. 2014, 8, 293. 10.3389/fncel.2014.00293.

(40) Yu, Y.; Herman, P.; Rothman, D. L.; Agarwal, D.; Hyder, F. Evaluating the Gray and White Matter Energy Budgets of Human Brain Function. Journal of Cerebral Blood Flow & Metabolism 2018, 38 (8), 1339–1353. 10.1177/0271678X17708691.

(41) Naundorf, B.; Wolf, F.; Volgushev, M. Unique Features of Action Potential Initiation in Cortical Neurons. Nature 2006, 440 (7087), 1060–1063. 10.1038/nature04610.

(42) Ramos-Brossier, M.; Romeo-Guitart, D.; Lanté, F.; Boitez, V.; Mailliet, F.; Saha, S.; Rivagorda, M.; Siopi, E.; Nemazanyy, I.; Leroy, C.; Moriceau, S.; Beck-Cormier, S.; Codogno, P.; Buisson, A.; Beck, L.; Friedlander, G.; Oury, F. Slc20a1 and Slc20a2 Regulate Neuronal Plasticity and Cognition Independently of Their Phosphate Transport Ability. Cell Death Dis. 2024, 15 (1), 20. 10.1038/s41419-023-06292-z.

(43) Maljevic, S.; Lerche, H. Potassium Channels: A Review of Broadening Therapeutic Possibilities for Neurological Diseases. J. Neurol. 2013, 260 (9), 2201–2211. 10.1007/s00415-012-6727-8.

(44) Catela, C.; Chen, Y.; Weng, Y.; Wen, K.; Kratsios, P. Control of Spinal Motor Neuron Terminal Differentiation through Sustained Hoxc8 Gene Activity. Elife 2022, 11. 10.7554/eLife.70766.

(45) Kim, M.; Fontelonga, T. M.; Lee, C. H.; Barnum, S. J.; Mastick, G. S. Motor Axons Are Guided to Exit Points in the Spinal Cord by Slit and Netrin Signals. Dev. Biol. 2017, 432 (1), 178–191. 10.1016/j.ydbio.2017.09.038.

(46) Lim, S. H.; Sung, Y. J.; Jo, N.; Lee, N. Y.; Kim, K. S.; Lee, D. Y.; Kim, N. S.; Lee, J.; Byun, J. Y.; Shin, Y. B.; Lee, J. R. Nanoplasmonic Immunosensor for the Detection of SCG2, a Candidate Serum Biomarker for the Early Diagnosis of Neurodevelopmental Disorder. Sci. Rep. 2021, 11 (1), 1–11. 10.1038/s41598-021-02262-7.

(47) Vidal, O. M.; Vélez, J. I.; Arcos-Burgos, M. ADGRL3 Genomic Variation Implicated in Neurogenesis and ADHD Links Functional Effects to the Incretin Polypeptide GIP. Sci. Rep. 2022, 12 (1), 1–14. 10.1038/s41598-022-20343-z.

(48) Rasband, M. N. The Axon Initial Segment and the Maintenance of Neuronal Polarity. Nat. Rev. Neurosci. 2010, 11 (8), 552–562. 10.1038/nrn2852.

(49) Chen, C.; Jonas, P. Synaptotagmins: That’s Why So Many. Neuron 2017, 94 (4), 694–696. 10.1016/j.neuron.2017.05.011.

(50) Liu, Q.; Xie, F.; Siedlak, S. L.; Nunomura, A.; Honda, K.; Moreira, P. I.; Zhua, X.; Smith, M. A.; Perry, G. Neurofilament Proteins in Neurodegenerative Diseases. Cellular and Molecular Life Sciences 2004, 61 (24), 3057–3075. 10.1007/s00018-004-4268-8.

(51) Cohen, L. D.; Zuchman, R.; Sorokina, O.; Müller, A.; Dieterich, D. C.; Armstrong, J. D.; Ziv, T.; Ziv, N. E. Metabolic Turnover of Synaptic Proteins: Kinetics, Interdependencies and Implications for Synaptic Maintenance. PLoS One 2013, 8 (5), e63191. 10.1371/journal.pone.0063191.

(52) Chepyala, S. R.; Liu, X.; Yang, K.; Wu, Z.; Breuer, A. M.; Cho, J. H.; Li, Y.; Mancieri, A.; Jiao, Y.; Zhang, H.; Peng, J. JUMPt: Comprehensive Protein Turnover Modeling of in Vivo Pulse SILAC Data by Ordinary Differential Equations. Anal. Chem. 2021, 93 (40), 13495–13504. 10.1021/acs.analchem.1c02309.

(53) Wiśniewski, J. R.; Hein, M. Y.; Cox, J.; Mann, M. A “Proteomic Ruler” for Protein Copy Number and Concentration Estimation without Spike-in Standards. Molecular and Cellular Proteomics 2014, 13 (12), 3497–3506. 10.1074/mcp.M113.037309.

