## Supplemental Information for "A Deep Quantitative Proteome Turnover Platform for Human iPSC-derived Neurons"

### Table of Contents


- **Supplemental Methods**: Human iPSC-derived Neuron Culture.
- **Supplementary Figure S1:** Correlation analyses of DDA and DIA dynamic SILAC proteomics data from human iPSC-derived neurons.
- **Supplementary Figure S2:** Comparing protein half-lives from different protein classes in human motor and cortical neurons.
- **Supplementary Figure S3:** GO-enrichment analyses showing enriched molecular functions of proteins with different ranges of half-lives in human neurons.
- **Supplementary Figure S4:** Heatmap clustering of synaptic protein half-lives measured in our human neuron data and published datasets from rodent neurons and brain tissues.
- **Supplementary Table S1**: Neuron specific contaminant library.
- **Supplementary Table S2**: Protein turnover and abundance in human iPSC-derived cortical and motor neurons.

#### Supplemental Methods:

##### Human iPSC-derived Neuron Culture

Human iPSCs were maintained in Essential 8 medium (Gibco) in 10 cm dishes coated with Matrigel (Corning). Human iPSCs were differentiated into glutamatergic cortical neurons and spinal motor neurons under two different protocols. Glutamatergic cortical neurons were produced using i$^3$Neuron technology in two weeks.[1–3] Briefly, wild-type human iPSCs with stably integrated doxycycline-inducible neurogenin2 (NGN2) cassette were dissociated with Accutase and plated into neuron induction medium (2 µg/mL of doxycycline, 10 µM of ROCK inhibitor Y-27632, N2 supplement, non-essential amino acid (NEAA) supplement, GlutaMAX, and DMEM/F12 with HEPES) on Matrigel-coated plates for 3 days with daily medium changess. Then day-3 neurons were dissociated with Accutase and replated onto poly-L-ornithine (PLO) coated plates in neuron medium (DMEM/F12 for SILAC medium supplemented with heavy or light arginine and lysine, N2 supplement, B-27 supplement, NEAA, 10 ng/mL of BDNF, 10 ng/mL of GDNF, 10 ng/mL of NT-3, 0.2 µg/mL of Laminin, 0.2 µg/mL of doxycycline and GlutaMAX). Cortical neurons were maintained with warm half-medium change every other day until maturation at 2 weeks.

Spinal motor neurons were produced using a series of small molecule cocktails that guided the iPSC through development and maturation over the course of 28 days.[4] To start the differentiation, iPSC medium was supplemented with 3 µM of CHIR99021, 2 µM of DMH-1, and 2 µM of SB4431542 for 6 days to induce the formation of neuroepithelial progenitor cells. Then neuroepithelial progenitor cells were replated into medium supplemented with 0.1 µM of retinoic acid, 0.5 µM of purmophamine, 1 µM of CHIR99021, 2 µM of DMH-1, and 2 µM of SB4431542 for 6 days. Immature motor neurons were obtained by culturing progenitor cells in neural medium with 0.5 µM of retinoic acid and 1 µM of Pur for 8 days. Postmitotic neurons were obtained by supplementing the neural medium with 0.5 μM of retinoic acid, 1 μM of Pur, and 0.1 μM of Compound E for 10 days. Warm half-medium changes were conducted every other day.

**References:**


(1) Wang, C.; Ward, M. E.; Chen, R.; Liu, K.; Tracy, T. E.; Chen, X.; Xie, M.; Sohn, P. D.; Ludwig, C.; Meyer-Franke, A.; Karch, C. M.; Ding, S.; Gan, L. Scalable Production of IPSC-Derived Human Neurons to Identify Tau-Lowering Compounds by High-Content Screening. *Stem Cell Reports* **2017**, *9* (4), 1221–1233. https://doi.org/10.1016/j.stemcr.2017.08.019.

(2) Fernandopulle, M. S.; Prestil, R.; Grunseich, C.; Wang, C.; Gan, L.; Ward, M. E. Transcription Factor–Mediated Differentiation of Human IPSCs into Neurons. *Curr Protoc Cell Biol* **2018**, *79* (1), 1–48. https://doi.org/10.1002/cpcb.51.

(3) Hasan, S.; Fernandopulle, M. S.; Humble, S. W.; Frankenfield, A. M.; Li, H.; Prestil, R.; Johnson, K. R.; Ryan, B. J.; Wade-Martins, R.; Ward, M. E.; Hao, L. Multi-Modal Proteomic Characterization of Lysosomal Function and Proteostasis in Progranulin-Deficient Neurons. *Mol Neurodegener* **2023**, *18* (1). https://doi.org/10.1186/s13024-023-00673-w.

(4) Du, Z. W.; Chen, H.; Liu, H.; Lu, J.; Qian, K.; Huang, C. T. L.; Zhong, X.; Fan, F.; Zhang, S. C. Generation and Expansion of Highly Pure Motor Neuron Progenitors from Human Pluripotent Stem Cells. *Nat Commun* **2015**, *6*, 1–9. https://doi.org/10.1038/ncomms7626.

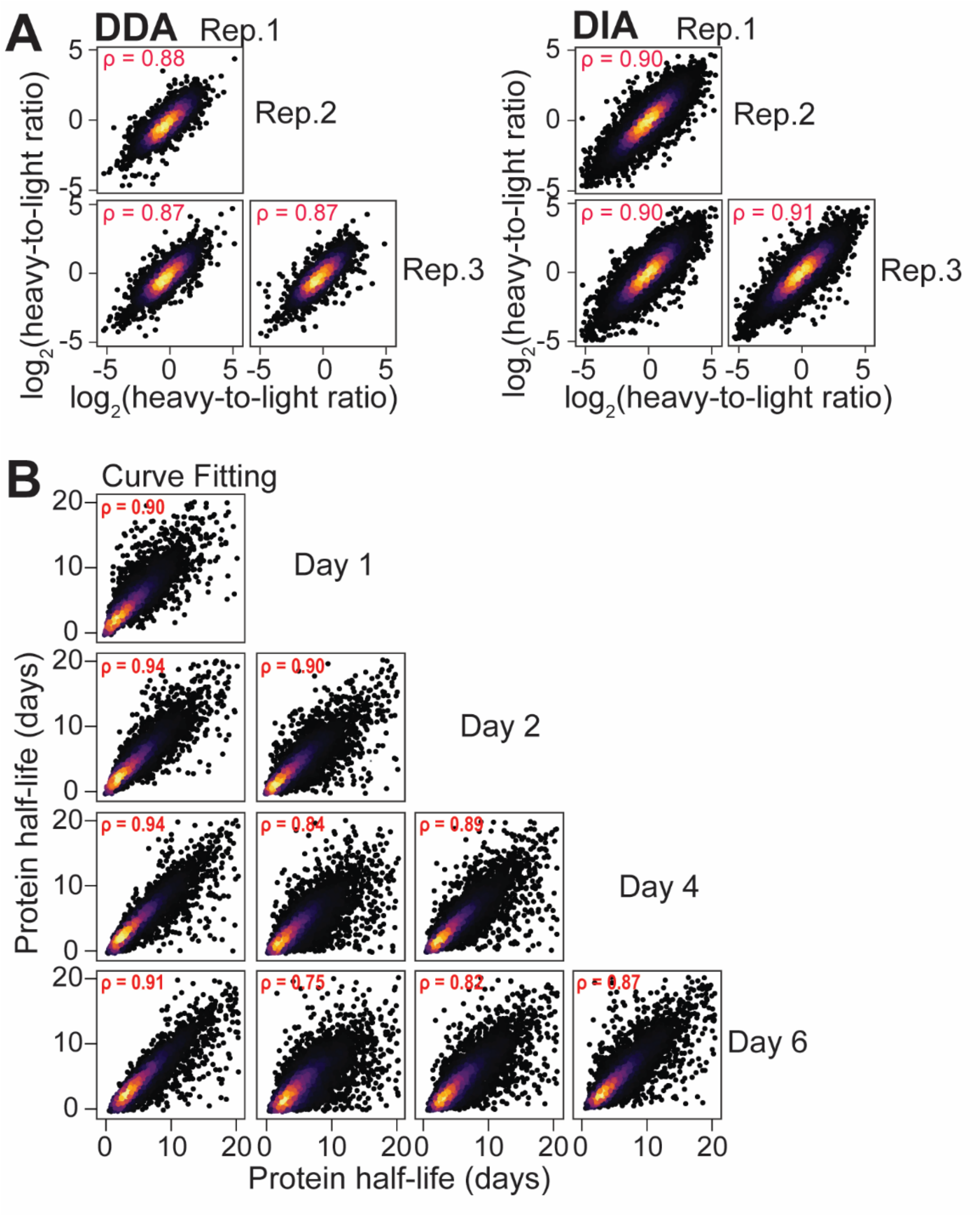


**Supplementary Figure S1: Correlation analyses of DDA and DIA dynamic SILAC proteomics data from human iPSC-derived neurons.** (A) Correlations of technical replicates in DDA and DIA dynamic SILAC proteomics data. ρ indicates Spearman's correlation. (B) Correlations of protein half-lives measured by multiple time point curve fitting and single time point equations.

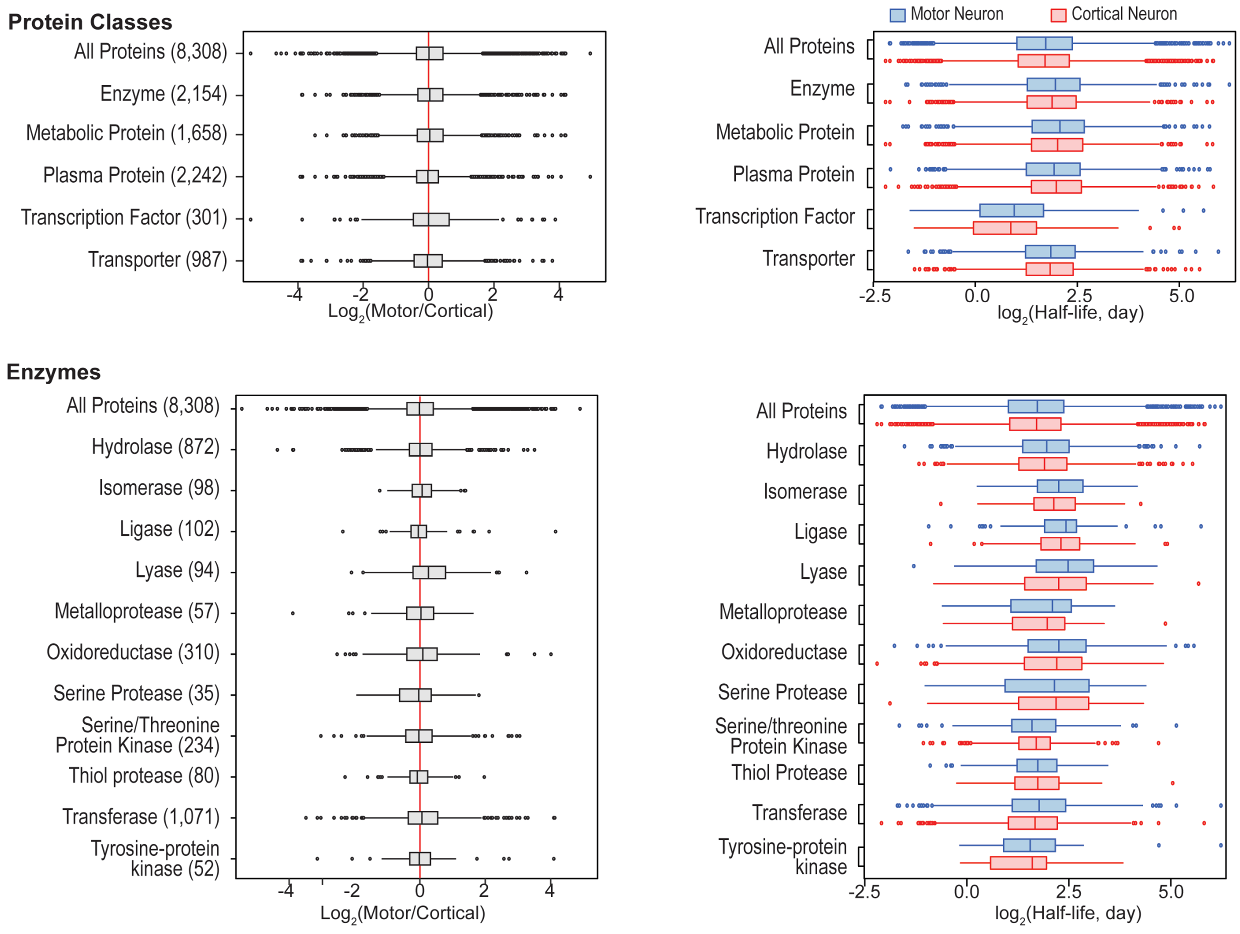


**Supplementary Figure S2: Comparing protein half-lives from different protein classes in human motor and cortical neurons.** Boxplots comparing the fold change (left) and distribution (right) of protein half-lives quantified in motor and cortical neurons. The number of proteins in each category is indicated in parentheses.

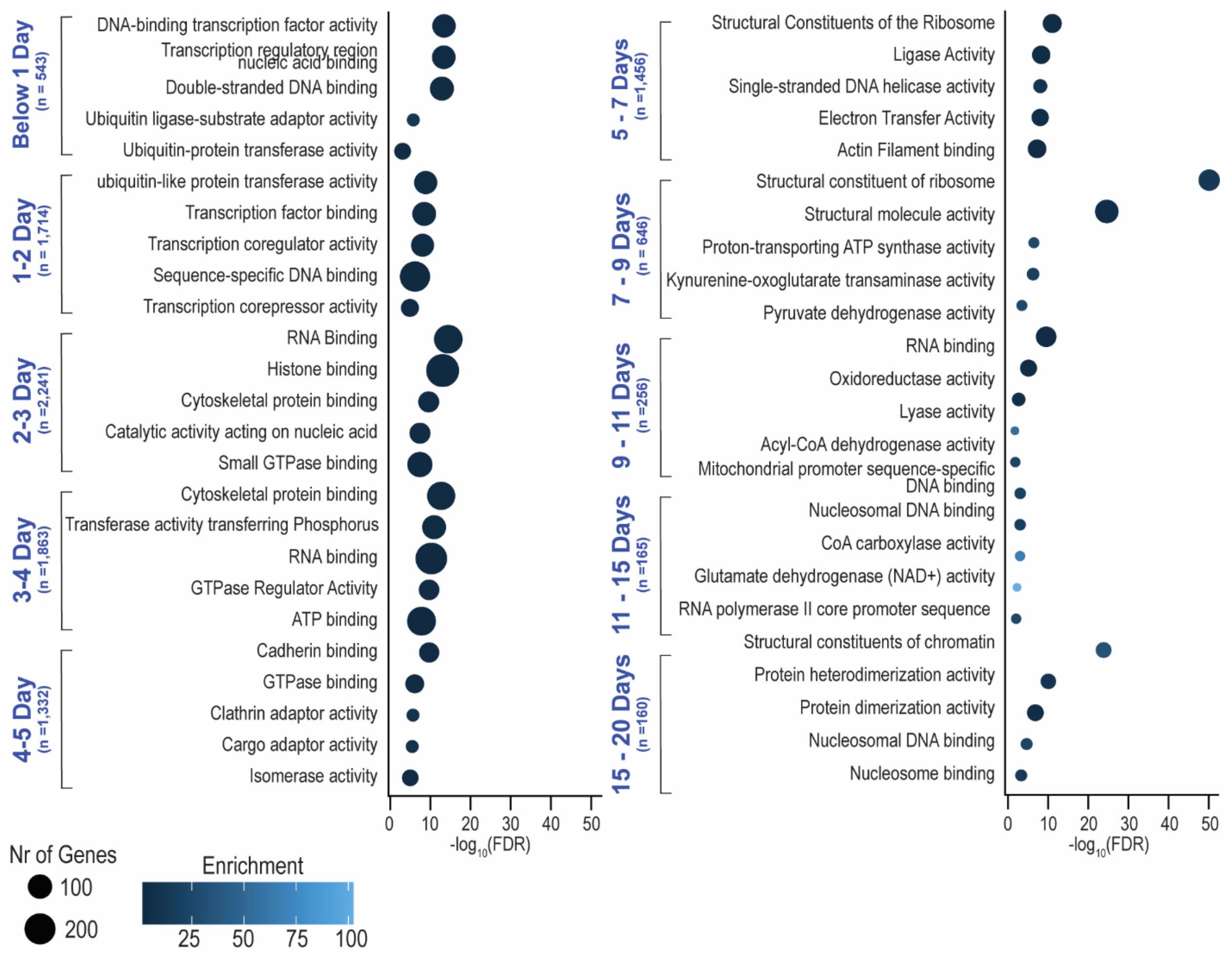


**Supplementary Figure S3: GO-enrichment analyses showing enriched molecular functions of proteins with different ranges of half-lives in human neurons.**

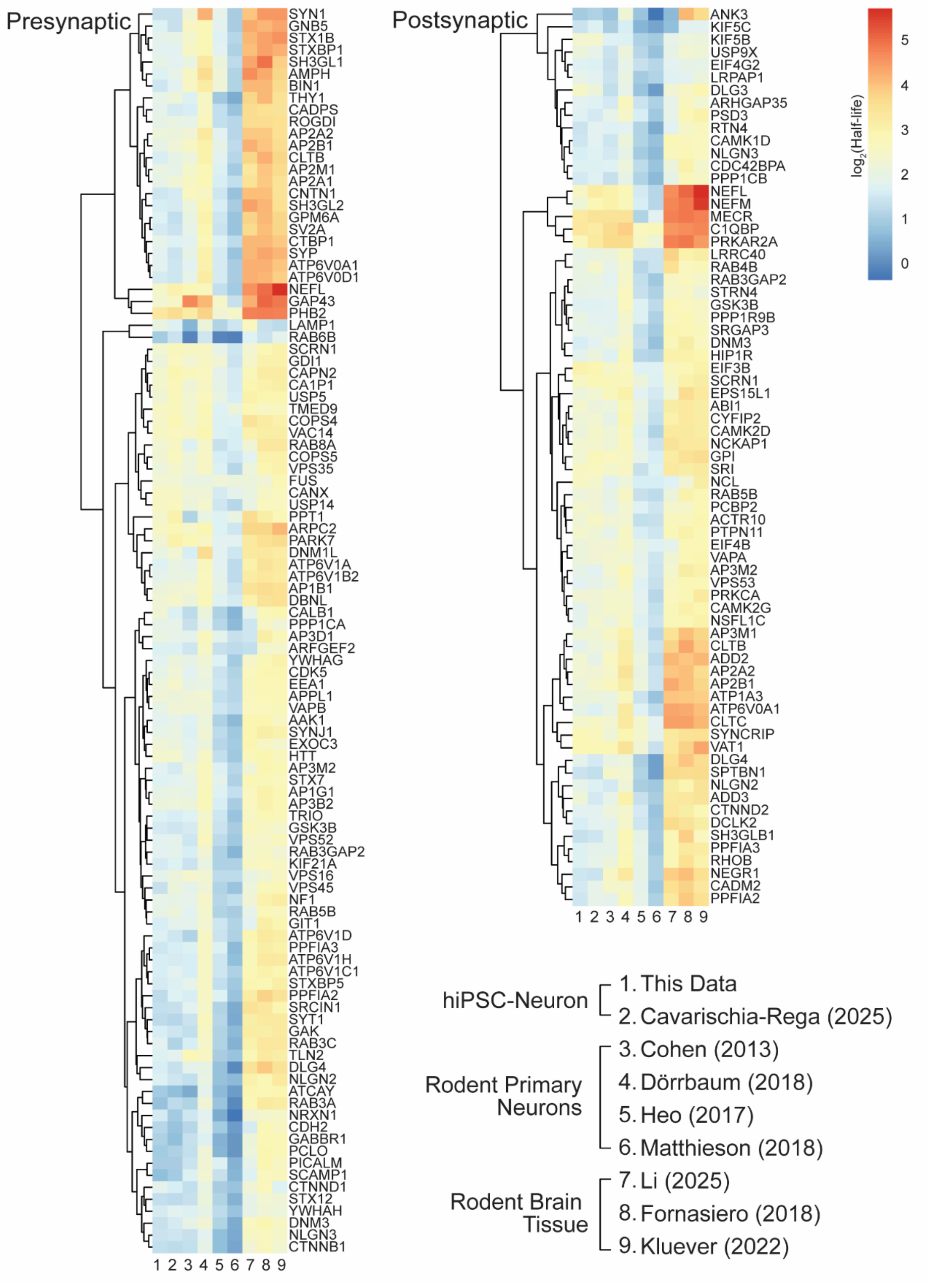


**Supplementary Figure S4: Heatmap clustering of synaptic protein half-lives measured in our human neuron data and published datasets from rodent neurons and brain tissues.**
